# The potential applications of high-resolution 3D scanners in the taxonomic classification of insects

**DOI:** 10.1101/2024.06.17.599367

**Authors:** Cameron J Peacock, William Romeu-Evans, Christopher Hassall, Simon Goodman

## Abstract

The phenotypic classification of small biological specimens, such as insects, can be dependent on phenotypic features that are difficult to observe and communicate to others. Here, we evaluate how high-resolution 3D photogrammetric scanner technology can potentially allow such features to be resolved and visualised as a 3D models, which can then be shared as a taxonomical resource for species identification, as ‘virtual type specimens’, and for educational and public engagement purposes. We test the viability and limitations of this approach using specimens digitised with a Artec Micro scanner. Ten samples from unique species were mounted and scanned. The model outputs were evaluated against an identification key, which compiled diagnostic features for the specimens from the wider literature, to describe the specimens to the lowest taxonomic level possible. The results showed that six of the ten specimens could be identified to species level using the scans. Threshold values for body length and width were 10.7 mm and 4.4 mm respectively. Below these body dimensions important diagnostic features of specimens could not be resolved reliably. This result suggests that with current technology, 3D photogrammetric modelling is a viable method for taxonomic identification of a wide range of insect groups with larger body sizes. This approach opens up novel applications for species identification and data sharing among taxonomists, international field research, conservation efforts, and entomological outreach. However, the limitations of this approach to taxonomic identification must be considered depending upon the size of the specimen and its diagnostic features. Future developments in the technology and processing methods used may alleviate the constraints on body size exhibited in this study, widening the applications for smaller bodied specimens.

## Introduction

The current number of documented insect species is estimated at over one million, suggesting that only 15 to 20 percent of insect species have been classified (Foottit and Adler, 2009). Insects play a pivotal role in global ecosystems, acting as food sources (Tallamy and Shriver, 2021), mediating ecosystem services such as pollination and nutrient recycling (Schowalter *et al*., 2018), as well as some being disease vectors and parasites (Lounibos, 2002). The sheer diversity of insects has required the development of extensive and detailed species identification and description techniques, with an emphasis on rapid and effective identification. Despite the importance of taxonomic tools and skills to biodiversity research, concerns have been raised regarding the future of species identification due to loss of taxonomic expertise as experienced researchers retire, and low training rates of new specialists, resulting in a decreasing capacity for species identification (Lee, 2000). Various pressures such as climate change and human activity are exacerbating these concerns by driving many species towards extinction at a rapid rate, creating a biodiversity crisis which requires extensive monitoring (Bellard *et al*., 2012).

Three-dimensional photogrammetric scanning offers a promising enhancement to contemporary taxonomic practice. Photogrammetric techniques generate 3D models of scanned objects and present a range of benefits over current high-resolution methods used for taxonomic identification (e.g., SEM and Micro-CT scanners; see Table 1 for a detailed comparison). These include its low cost, its ability to rapidly generate high-fidelity 3D models with little preparation requirements and its user-friendly programming requiring little training. Another key benefit of this approach is the ability to share generated 3D models digitally, reducing the need for specimen preservation and transportation. This reduces costs further and streamlines the identification process, with potential to help mitigate some of the issues around erosion of taxonomic expertise by facilitating access to specimens and training, and by opening new possibilities for artificial intelligence supported taxonomic identification. This technology has been applied to various areas of taxonomic research, including virtual palaeontology for fossil reconstruction (Bartolini-Lucenti *et al*., 2021; Theodorou *et al*., 2018), detailed identification and cataloguing of arthropod species (Hita-Garcia *et al*, 2017), and in institutions that act as reference archives, removing the need for physical specimens through digitisation in natural history collections (Plum and Labonte, 2021; Rybenská and Borůvková, 2020). Here, we evaluate the practical taxonomic ability of photogrammetric 3D scanning technology to resolve characters important for taxonomic identification of insects with varying sizes and surface features. The novelty of this research comes from the direct measurement of taxonomic utility of this approach under practical conditions, which provides empirical data to assess the limitations of current 3D scanning technology.

**Table 1.**
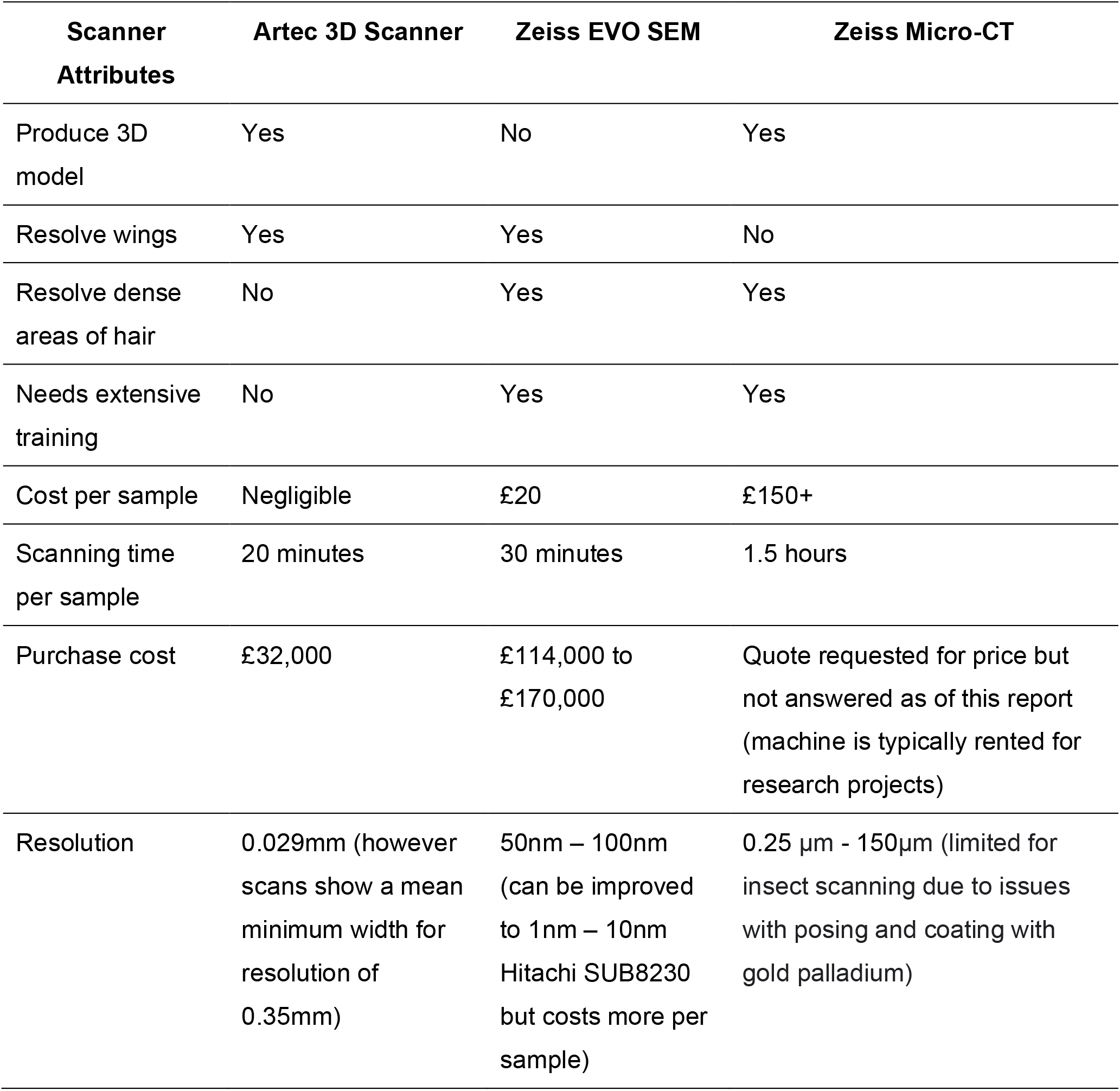
A comparison of scanner attributes between the Artec 3D Micro Scanner, Zeiss EVO SEM and Zeiss Micro-CT. Information sourced from personal communication (see supplementary material).

| Scanner Attributes | Artec 3D Scanner | Zeiss EVO SEM | Zeiss Micro-CT |
| --- | --- | --- | --- |
| Produce 3D model | Yes | No | Yes |
| Resolve wings | Yes | Yes | No |
| Resolve dense areas of hair | No | Yes | Yes |
| Needs extensive training | No | Yes | Yes |
| Cost per sample | Negligible | £20 | £150+ |
| Scanning time per sample | 20 minutes | 30 minutes | 1.5 hours |
| Purchase cost | £32,000 | £114,000 to £170,000 | Quote requested for price but not answered as of this report (machine is typically rented for research projects) |
| Resolution | 0.029mm (however scans show a mean minimum width for resolution of 0.35mm) | 50nm – 100nm (can be improved to 1nm – 10nm Hitachi SUB8230 but costs more per sample) | 0.25 µm - 150µm (limited for insect scanning due to issues with posing and coating with gold palladium) |

## Methods

We examined 10 specimens from different species, which were selected to give a range of different body sizes and morphological characteristics. Specimens were scanned using an Artec Micro scanner (Artec3D, Luxembourg) with a precision of 10µm and a resolution of 29µm. The individuals chosen represented species from Coleoptera, Lepidoptera, Diptera and Orthoptera (Table 2). Specimens ranged in body length from 1.63 mm to 33.49 mm and possessed unique diagnostic traits to allow for identification, such as wing morphology, body colour and pattern etc. Bat fly (Nycteribiidae spp.) and bat bug specimens (Cimidae spp) (n=2), originally sampled in Baja California, Mexico (Najera-Cortazar et 2023), were obtained from a collection held at the University of Leeds. The remaining specimens were sourced from a routine insect monitoring programme operating at Hatfield Moors, UK (53.533°N, -0.978°E). Samples had been preserved in 70% ethanol and therefore required rehydration using acetone and a heat lamp (Grissell and Schauff, 1995). Based on the scanner manufacturers recommendations, each specimen was then sprayed with a thin layer of Aesub blue scanning spray to create a matte finish and prevent glare during scanning. Each specimen was mounted on a needle and secured on the scanning plate. The small complex fine setting was selected on the Artec Studio 3D software and once initiated the scanning arm rotated around the sample, taking 30 scans from a range of angles. These were the default settings, however additional scans could be subsequently added. The scans are automatically compiled into a 3D model for each specimen which was then refined using the sharp fusion and hole filling tools in the Artec Studio program to generate a watertight model (a complete model with no unfilled areas or holes). The models were then used to determine body length, width, wing length and volume for each specimen.

**Table 2.** The identification key used for each specimen, including the significant diagnostic features used in the wider literature to identify various taxonomic groups. The sources of these diagnostic features can be found in Appendix 1. The table also includes the taxonomic score achieved for each specimen, with 0 meaning unidentifiable and 1 to 5 corresponding to class, order, family, genus, and species respectively. The body length of each species is also provided.

| Scientific Name | Identification Characters | Taxa score<br>(0-5) | Body Length<br>(mm) |
| --- | --- | --- | --- |
| Nycteribiidae spp. | None of the diagnostic features in the literature for identification, such as flattened body, lack of wings and ocelli and leg morphology, could be resolved. | 0 | 1.63 |
| Cimidae spp. | None of the diagnostic features described in the literature for identification, such as antennae length/ segment number, tarsi morphology, head shape and ocelli presence, length of fringe hairs on pronotum, could be resolved. | 0 | 3.70 |
| *Sarcophaga* spp. | 1. Jointed legs, hard cuticle, six legs, head/thorax/abdomen – **Insecta**<br>2. Has fully developed wings, one pair of well-developed wings with hindwings reduced to small knobs behind the forewings – **Diptera**<br>3. Not metallic and shiny, typically black, and grey. Dorsum of thorax often striped and abdomen chequered – **Sarcophagidae**<br>4. Terminal abdominal segment usually red, less than 10mm long, three dark stripes on thorax, large red eyes- **Sarcophaga**<br>5. Species are often difficult to distinguish, with the most reliable method involving examination of the genitalia under microscope. | 1 | 6.77 |
| *Coccinella septempunctata* | 1. Jointed legs, hard cuticle, six legs, head/thorax/abdomen – **Insecta**<br>2. Fully developed wings, visible mouthparts, forewings are shield-like and lack visible veins, forming a straight line where they meet dorsally – **Coleoptera** | 5 | 7.66 |
|  | 3. 1- 7mm in Length, body typically ovate or hemispherical, moderately to strongly convex, often with spots on pronotum and elytra – **Coccinellidae**<br>4/5. Orange/ red elytra with black spots, seven spots in total (key feature) – **Coccinella septempunctata** |  |  |
| *Phosphuga atrata* | 1. Jointed legs, hard cuticle, six legs, head/thorax/abdomen – **Insecta**<br>2. Fully developed wings, visible mouthparts, forewings are shield-like and lack visible veins, forming a straight line where they meet dorsally – **Coleoptera**<br>3. Relatively large (9-30mm), antennae widely spaced and insert on the lateral side of the head, flattened dorsally, tarsi of all six legs have 5 segments, antennae clubbed or wider at the tip – **Silphidae**<br>4/5. Uniform colouration (mainly dark), antennae expanded apically but lack clubbed tip, head elongated, elytra with raised longitudinal lines – **Phosphuga atrata** | 5 | 11.26 |
| *Tetrix undulata* | 1. Jointed legs, hard cuticle, six legs, head/thorax/abdomen – **Insecta**<br>2. Two pairs of developed wings, visible and developed mandible mouthparts, abdomen tip lacks obvious pincers, forewings leathery and veins are visible, body is not flattened and head visible from above, front legs are simple but hind legs are enlarged for jumping – **Orthoptera**<br>3. Antennae shorter than body, size between 8 and 14 mm, elongated pronotum, brown colouration - **Tetrigidae**<br>4/5. Pronotum not extending as far as hind knees, strong midline ridge, hind wings shorter than pronotum – **Tetrix undulata** | 5 | 13.29 |
| *Quedius xanthopus* | 1. Jointed legs, hard cuticle, six legs, head/thorax/abdomen – **Insecta**<br>2. Fully developed wings, visible mouthparts, forewings are shield-like and lack visible veins, forming a straight line where they meet dorsally – **Coleoptera**<br>3. 1 to 24mm in length, number of tarsi can be 3 for all six legs or 5 for all six legs, mainly characterised by short elytra, generally elongated and slender, abdomen is exposed – **Staphylinidae** | 3 | 14.66 |
|  | 4. Segments 1 and 2 of tarsi are broader than long, antennae are filiform (evenly curved segments give an even outline), elytra uniformly punctured – **Quedius** |  |  |
| *Eristalis pertinax* | 1. Jointed legs, hard cuticle, six legs, head/thorax/abdomen – **Insecta**<br>2. Has fully developed wings, one pair of well-developed wings with hindwings reduced to small knobs behind the forewings – **Diptera**<br>3. Typically short and segmented antennae, large eyes, presence of vena spuria (chitinized fold in the wing), some species mimic Hymenoptera but lack features such as stingers or two sets of wings – **Syrphidae**<br>4/5. Front tarsi are orange/yellow in colour, noticeable dark face stripe, curved rear tibia, 13 – 15mm in length, two vertical bands of hairs on the eyes – **Eristalis pertinax** | 5 | 15.41 |
| *Nicrophorus interruptus* | 1. Jointed legs, hard cuticle, six legs, head/thorax/abdomen – **Insecta**<br>2. Fully developed wings, visible mouthparts, forewings are shield-like and lack visible veins, forming a straight line where they meet dorsally – **Coleoptera**<br>3. Relatively large (9-30mm), antennae widely spaced and insert on the lateral side of the head, flattened dorsally, tarsi of all six legs have 5 segments, antennae clubbed or wider at the tip – **Silphidae**<br>4. Antennae knob-like with four segments – **Nicrophorus**<br>5. Black with orange/brown markings on elytra, hind tibia straight, club of antenna largely orange, anterior orange elytral markings widely separated – **Nicrophorus interruptus** | 5 | 20.75 |
| *Deilephila elpenor* | 1. Jointed legs, hard cuticle, six legs, head/thorax/abdomen – **Insecta**<br>2. Two pairs of well-developed wings, tube-like mouthparts, scaled wings which are powder-like – **Lepidoptera**<br>3. Forewing length greater than 1.2cm, forewing long and slender, pointed apically, frenulum well developed, body not excessively hairy – **Sphingidae**<br>4/5. wingspan 30 – 60mm, colouration includes browns and pinks, with unique banding and stripes being the key identifying feature – **Deilephila elpenor** | 5 | 33.49 |

A taxonomic identification key was generated by compiling a range of diagnostic features collected from pre-established taxonomic keys (see supplementary Appendix 1). This key along with the 3D generated models were then used to determine the taxonomic level to which each of the 10 specimens could be identified. Each specimen was then designated a taxonomic score based on this. A score of zero meant unidentifiable and scores ranging from one to five represented class, order, family, genus and species respectively.

Body size measurements were plotted against taxonomic scores for each specimen to estimate resolution threshold value of each trait (Figure 1). For the Artec scanner to be an effective tool for taxonomic identification we suggest that family should be the minimum level of classification achievable, as below this point diagnostic features become more diverse and difficult to distinguish (e.g., hair presence/length, wing venation). Family-level identification is also the standard taxonomic resolution in freshwater biomonitoring. Therefore, the point where the fit line intercepts the taxonomic score of 3 was used to determine the threshold value.

**Figure 1.**
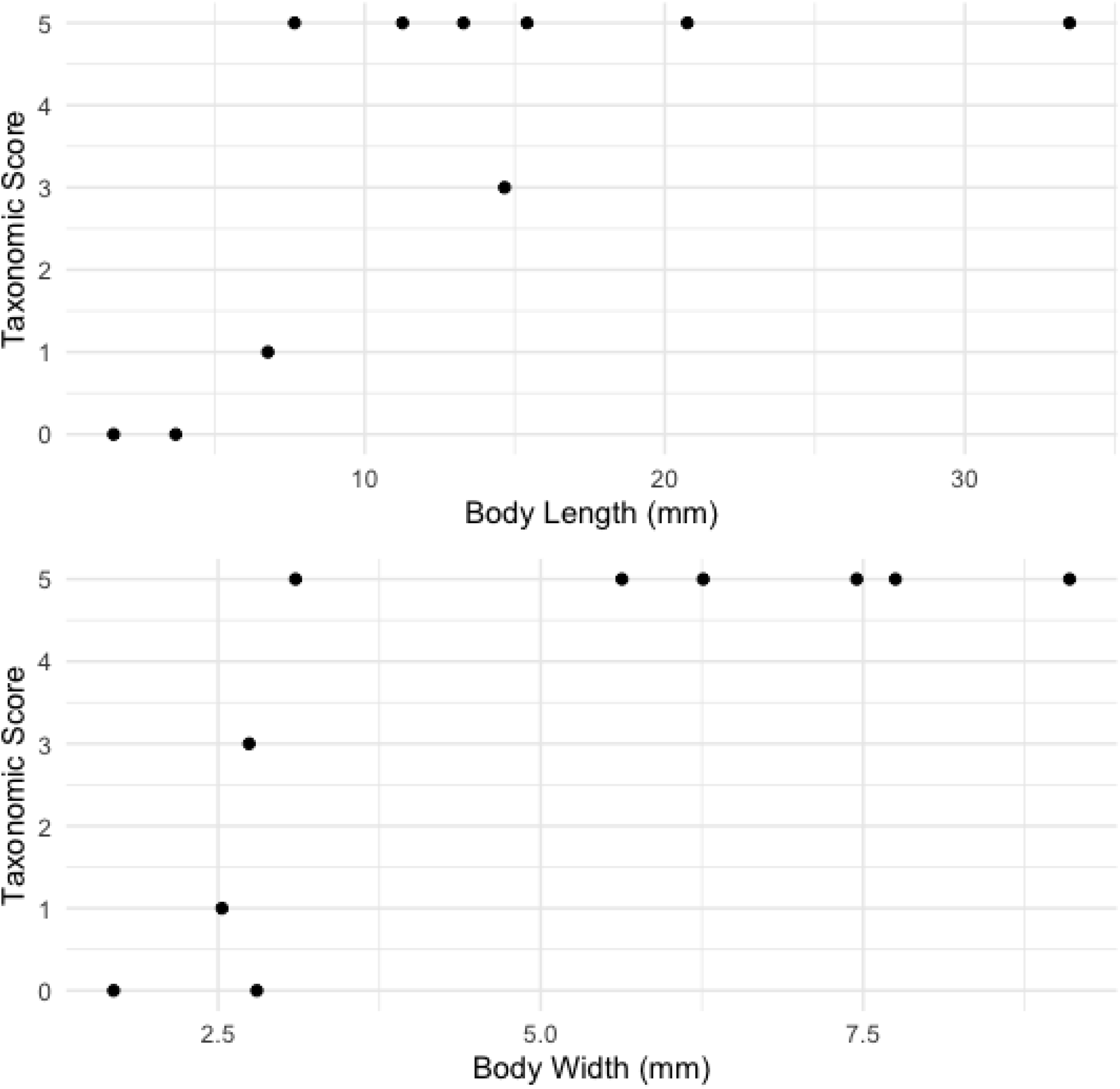
The relationship between two metrics of body size (graph A representing body length and graph B representing body width) and the taxonomic score achieved for each specimen. The taxonomic score is ranked from 0 to 5, with zero being unidentifiable and 1, 2, 3, 4 and 5 corresponding to class, order, family, genus, and species respectively.

## Results

It was found that seven of the total 10 specimens could be identified to family level or below, using diagnostic features compiled from numerous sources. The bat bug and bat fly ectoparasites, which had an average length of 2.67 mm, average width of 2.25 mm and average volume of 2.33 mm^3^ could not be identified to any taxonomic level. The third smallest specimen, *Sarcophaga* spp. could only be identified to the level of class (Insecta). This specimen had a body length of 6.77mm, body width of 2.53 mm and a volume of 29.39 mm^3^. The ranges for body size metrics for the remaining seven specimens were 7.66 mm to 33.49 mm for body length, 2.74mm to 9.10 mm for body width and 51.11 mm^3^ to 1394.22 mm^3^ for volume. Estimated resolution threshold values were 10.7 mm for body length, 4.4 mm for body width, 4.2 for surface area to volume ratio and 112 mm^3^ for volume. Finally, the minimum width resolved for each specimen was averaged, giving a mean of 0.35 mm, suggesting that any feature with a width lower than this are unlikely to be resolved. The process of using the 3D models to identify specimens to different taxonomic levels provided insight into the various diagnostic features which could be successfully resolved (Figure 2) and those that failed (Figure 3).

**Figure 2.**
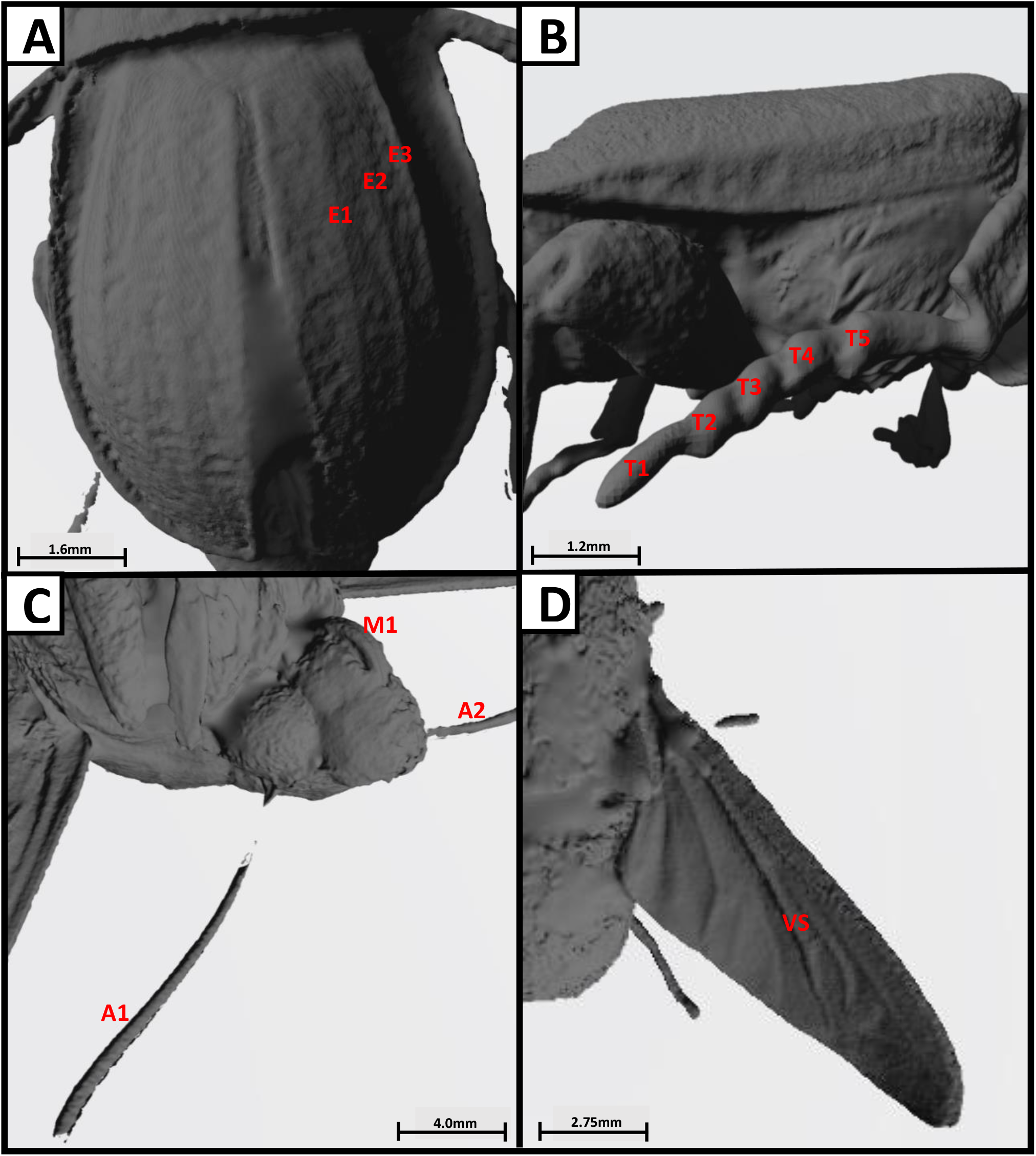
A collection of images taken of the Artec 3D scans of various specimens, highlighting various important diagnostic features for the identification of different taxonomic groups. A) depicts the raised longitudinal lines on the elytra of *Phosphuga atrata* (with three of these on the right side of the elytra being labelled E1, E2, E3). B) depicts the segments of the tarsi of *Nicrophorus interruptus* (labelled T1 to T5), used to identify *Nicophorus*. C) depicts the antennae and mouthpart of *Deilephila elpenor* (labelled M1 for mouthpart and A1/A2 for the antennae), used to identify Lepidoptera D) depicts the vena spuria (labelled VS) present in the centre of the wing of *Eristalis pertinax*, used to identify Syrphidae. Scale bar is attributed to each photo as scale varies between scan captures.

**Figure 3.**
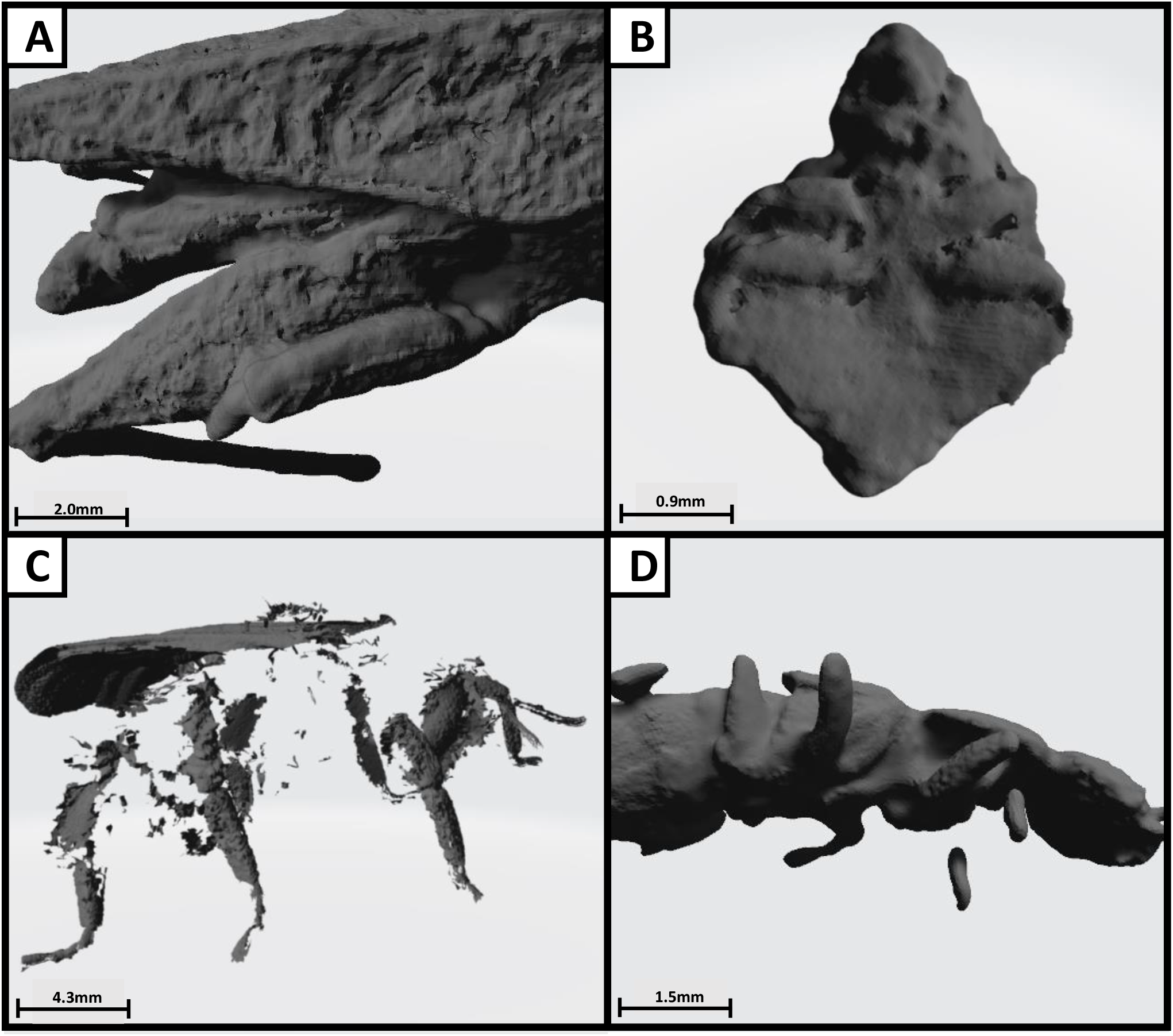
A collection of images taken of the Artec 3D scans of various specimens, highlighting limitations of this approach when trying to resolve various taxonomic groups. A) depicts the fusion of limbs of *Tetrix undulata*. B) depicts the inability to resolve features of a small bat bug specimen. C) depicts the inability to resolve areas of densely packed hair of a *Bombus* species. D) Depicts the incomplete resolution of the lower legs of *Quedius xanthopus*. Scale bar is attributed to each photo as scale varies between scan captures.

## Discussion

This study was carried out to evaluate the application of 3D scanning for taxonomic identification of insects and determine the threshold values for body size where resolution of important diagnostic features becomes limited. The Artec 3D scanner is capable of resolving a range of diagnostic features needed for taxonomic identification of species from a range of groups, including Lepidoptera, Diptera and Coleoptera (see Figure 2). This is significant due to the erosion of taxonomic expertise due to retirement of taxonomists and low training rates of new specialists, which is becoming a growing concern in the field of species identification and biodiversity management. This also highlights its potential future application to novel species identification. This study identified key limitations of using 3D modelling for arthropod taxonomic identification, particularly focusing on body size and the resolution of fine diagnostic features. Additional issues with this approach included fusing of limbs and inability to resolve areas of dense hairs (for example in *Bombus* spp.). Future research should aim to investigate the furthest limits of this technology, by including a wider range of species from various families. By increasing the number of unique individuals, a database of diagnostic features could be created to inform future applications of this technology to taxonomic identification. Further detail could be added using textures captured with high resolution photographs.

In comparison to other high-resolution imaging technologies, such as scanning electron microscopy and Micro-CT scanning, the Artec 3D scanner provides many benefits. These include the ability to manipulate the models’ orientation to allow for viewing of the specimen from various angles (360° viewing; Figure 4) and the ability to zoom into small diagnostic features. Additionally, the 3D photogrammetric scanners involve lower purchasing and running costs, shorter processing times and require less training than other conventional high resolution scanning techniques. Next steps would be blind trials with taxonomists, asking groups to identify specimen sets using 3D models without prior knowledge. This would help refine scanner protocols for taxonomic identification and demonstrate how this method can increase capacity for species identification. Additionally, recent developments of new 3D scanning technology such as scAnt, an open-source cost-effective platform, provide the opportunity for comparative empirical studies to examine the efficacy of different technologies in terms of taxonomic identification (Plum and Labonte, 2021). If these technologies are applied on a wider scale, this approach has the potential to enhance novel species discovery, support conservation efforts involving rare or undescribed species, and to promote public engagement around insect biodiversity topics.

**Figure 4.**
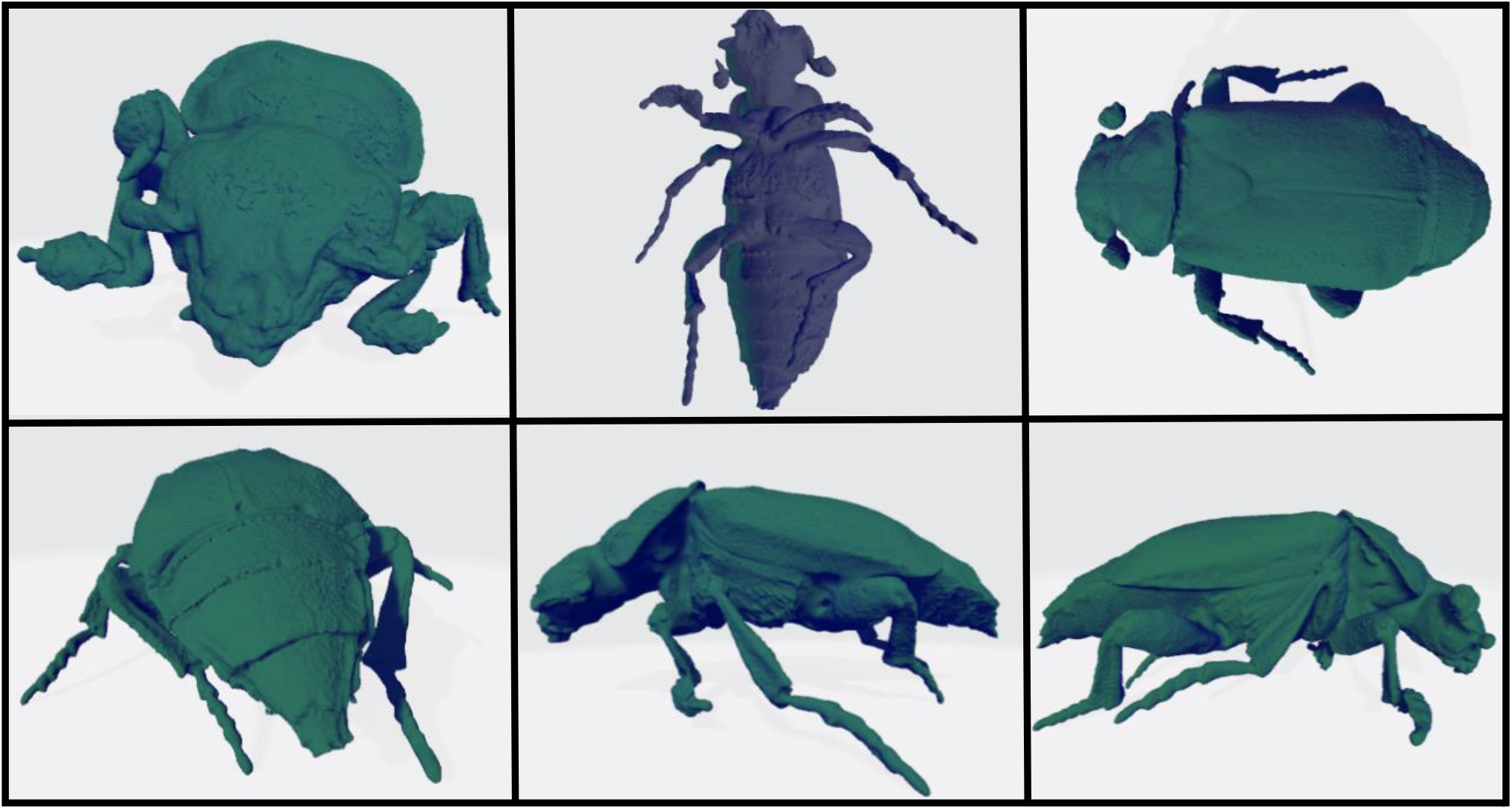
A collection of images taken of the Artec 3D scan of *Nicrophorus interruptus*, highlighting a subset of the various orientations that the model can be observed from. Viewing the model in the program Artec studio allows for a 360° view of the specimen, with the ability to zoom into important diagnostic features and manipulate the orientation of the model.

## Supporting information

Raw data

## Acknowledgements

The authors thank Dr Steve Compton for providing the specimens used in this study.

## Appendix 1

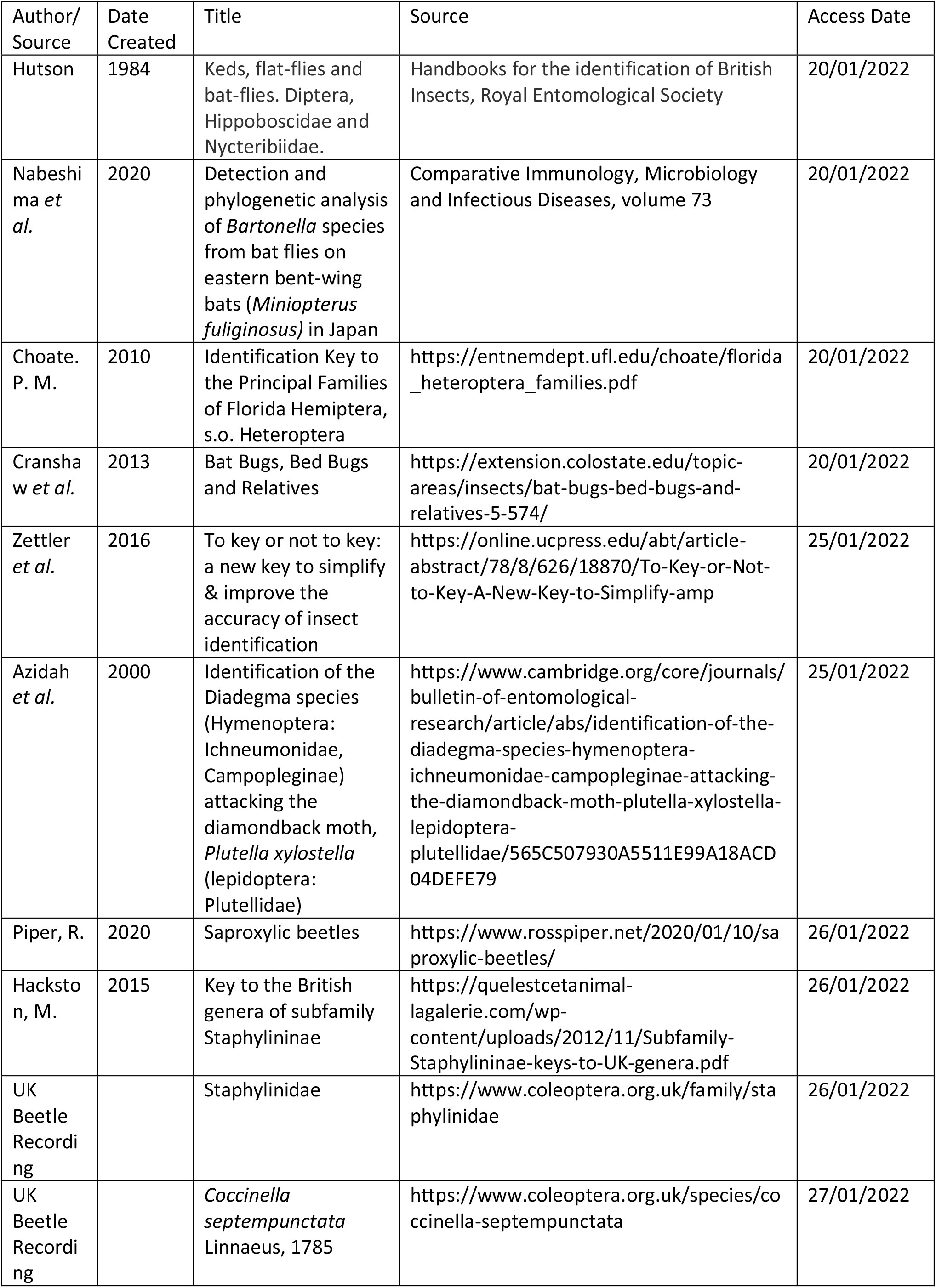
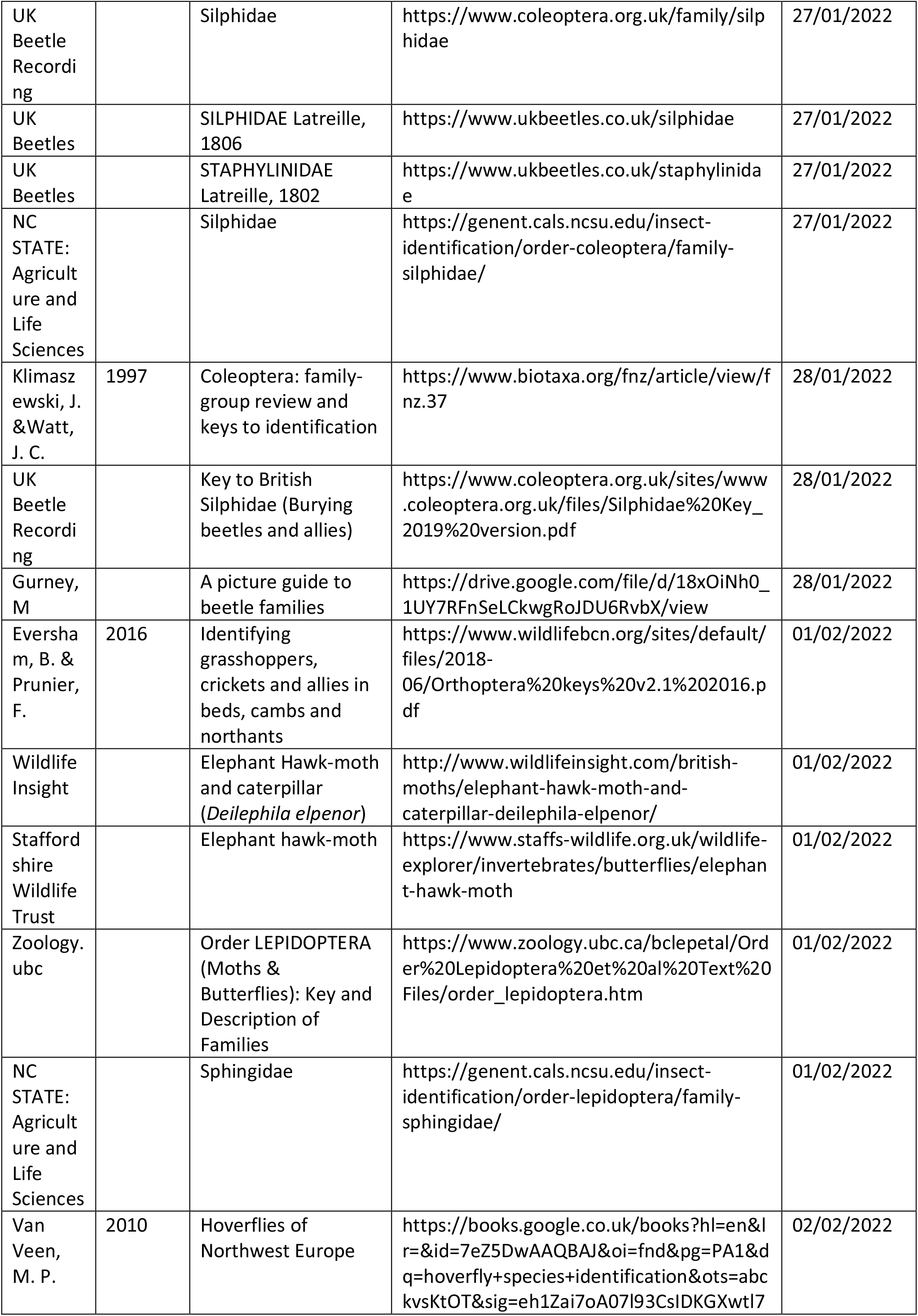
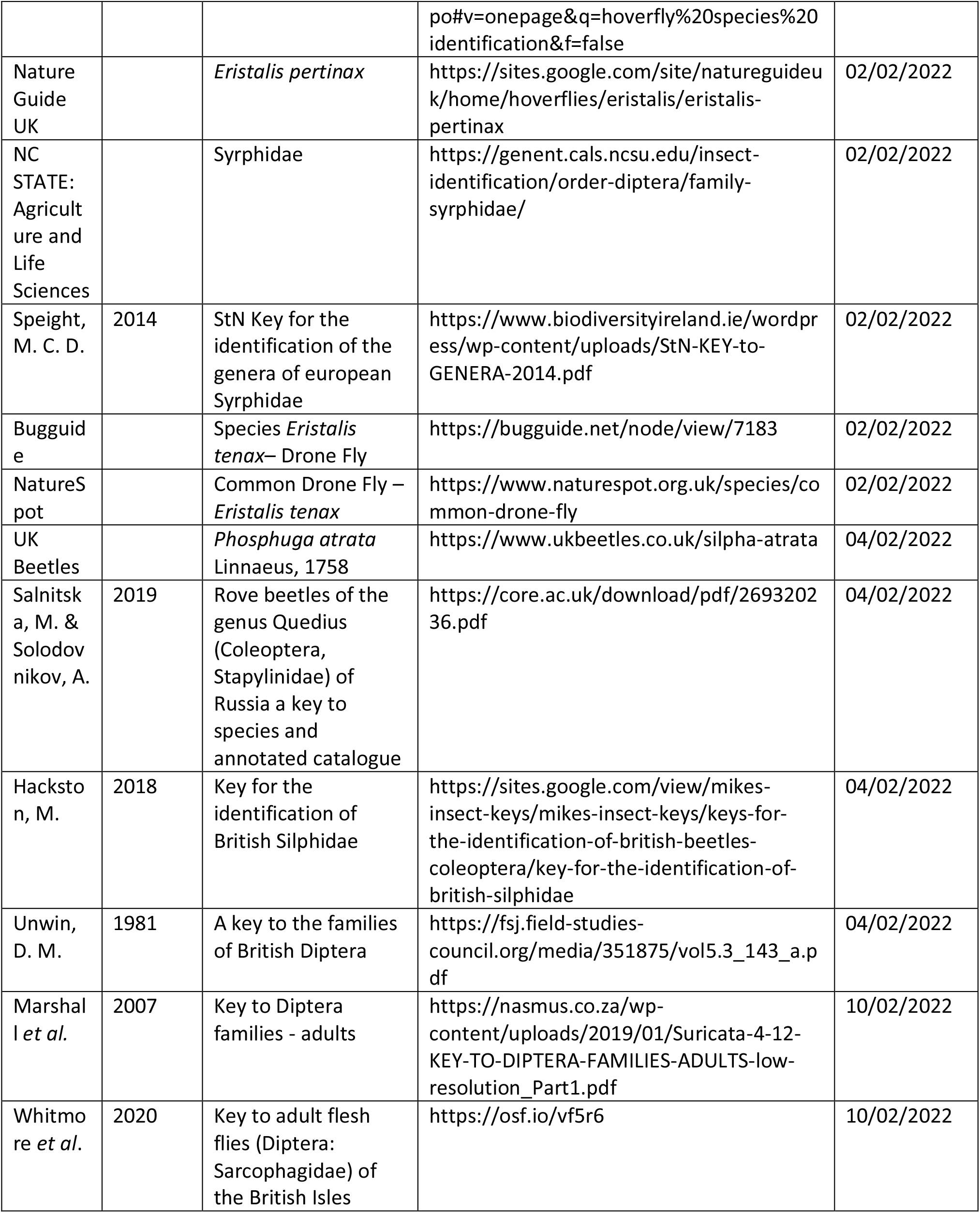

## Appendix 2

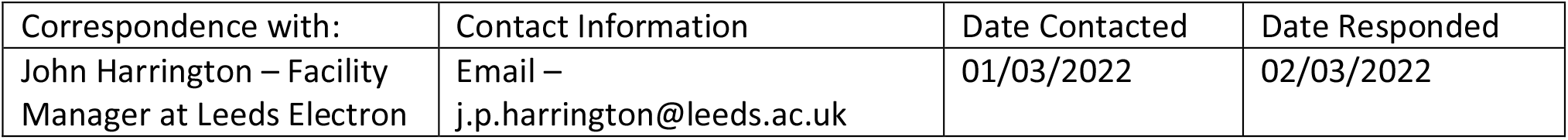
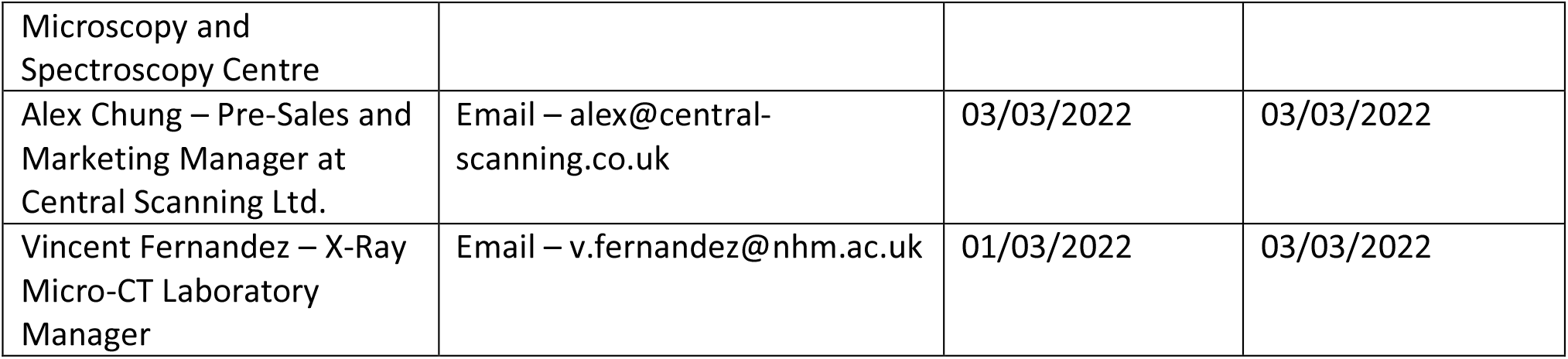

## References

Bartolini-Lucenti, S., Cirilli, O., Pandolfi, L., Sami, M., Dedola, G.L. and Rook, L. (2021). Applications of virtual paleontology on fossil remains and bone blocks of Montincino quarry (Brisighella, RA). La Fauna Messiniana di Cava Monticino, pp.37–48.

Bellard, C., Bertelsmeier, C., Leadley, P., Thuiller, W. and Courchamp, F. (2012). Impacts of climate change on the future of biodiversity. Ecology Letters, 15(4), pp.365–377.

Foottit, R.G. and Adler, P.H. (2009). Insect biodiversity: science and society. John Wiley & Sons, Ltd. pp.1–359.

Guerra-García, J.M., Espinosa Torre, F. and García Gómez, J.C. (2008). Trends in taxonomy today: an overview about the main topics in taxonomy. Zoológica Baetica, 19: 15–49.

Hita Garcia, F., Fischer, G., Liu, C., Audisio, T.L., Alpert, G.D., Fisher, B.L. and Economo, E.P. (2017). X-Ray microtomography for ant taxonomy: An exploration and case study with two new Terataner (Hymenoptera, Formicidae, Myrmicinae) species from Madagascar. PLoS One, 12(3), pp.1–36.

Lee, M.S. (2000). A worrying systematic decline. Trends in Ecology & Evolution, 15(8), pp.346.

Lounibos, L.P. (2002). Invasions by insect vectors of human disease. Annual Review of Entomology, 47(1), pp.233–266.

Najera-Cortazar, L. A., Keen, A., Kitching, T., Stokes, D., & Goodman, S. J. (2023). Phylogenetic analyses reveal bat communities in Northwestern Mexico harbor a high diversity of novel cryptic ectoparasite species. Ecology and Evolution, 13, e9645. 10.1002/ece3.9645

Plum, F. and Labonte, D. (2021). scAnt—an open-source platform for the creation of 3D models of arthropods (and other small objects). PeerJ, 9, p.e11155.

Rybenská, K. and Borůvková, B. (2020). Vybrané metody 3D digitalizace aplikované na příkladech hodinových exemplářů Náchodského muzea. Museum: Museum & Regional Studies, 58(2). pp.19–31.

Schowalter, T.D., Noriega, J.A. and Tscharntke, T. (2018). Insect effects on ecosystem services—Introduction. Basic and Applied Ecology, 26, pp.1–7.

Tallamy, D.W. and Shriver, W.G. (2021). Are declines in insects and insectivorous birds related?. The Condor, 123(1), pp.1–8.

Theodorou, G., Bassiakos, Y., Tsakalos, E., Yiannouli, E. and Maniatis, P. (2018). The use of CT scans and 3D modeling as a powerful tool to assist fossil vertebrate taxonomy. In Digital Heritage. Progress in Cultural Heritage: Documentation, Preservation, and Protection: 7th International Conference, EuroMed 2018, Nicosia, Cyprus, October 29–November 3, 2018, Proceedings, Part I 7. pp. 79–89.

